# Genome-Wide Association Study and transcriptome analysis reveals a complex gene network that regulates opsin gene expression and cell fate determination in *Drosophila* R7 photoreceptor cells

**DOI:** 10.1101/2024.08.05.606616

**Authors:** John C. Aldrich, Lauren A. Vanderlinden, Thomas L. Jacobsen, Cheyret Wood, Laura M. Saba, Steven G. Britt

## Abstract

**Background:** An animal’s ability to discriminate between differing wavelengths of light (i.e., color vision) is mediated, in part, by a subset of photoreceptor cells that express opsins with distinct absorption spectra. In *Drosophila* R7 photoreceptors, expression of the rhodopsin molecules, Rh3 or Rh4, is determined by a stochastic process mediated by the transcription factor *spineless*. The goal of this study was to identify additional factors that regulate R7 cell fate and opsin choice using a Genome Wide Association Study (GWAS) paired with transcriptome analysis via RNA-Seq.

**Results:** We examined Rh3 and Rh4 expression in a subset of fully-sequenced inbred strains from the *Drosophila* Genetic Reference Panel and performed a GWAS to identify 42 naturally-occurring polymorphisms—in proximity to 28 candidate genes—that significantly influence R7 opsin expression. Network analysis revealed multiple potential interactions between the associated candidate genes, *spineless* and its partners. GWAS candidates were further validated in a secondary RNAi screen which identified 12 lines that significantly reduce the proportion of Rh3 expressing R7 photoreceptors. Finally, using RNA-Seq, we demonstrated that all but four of the GWAS candidates are expressed in the pupal retina at a critical developmental time point and that five are among the 917 differentially expressed genes in *sevenless* mutants, which lack R7 cells.

**Conclusions:** Collectively, these results suggest that the relatively simple, binary cell fate decision underlying R7 opsin expression is modulated by a larger, more complex network of regulatory factors. Of particular interest are a subset of candidate genes with previously characterized neuronal functions including neurogenesis, neurodegeneration, photoreceptor development, axon growth and guidance, synaptogenesis, and synaptic function.

## Introduction

The initial steps of color vision are mediated by a population of specialized sensory neurons (i.e., photoreceptors) that express one or more spectrally-tuned photopigments called opsins (1–3). How an individual photoreceptor responds to light of a given wavelength is a function of the absorption spectra of the opsin it expresses in concert with various screening and/or activating pigments, while the overall diversity of photoreceptors within a retina determines the range and number of distinct colors that can be detected (4). Consequently, the mechanism through which an organism perceives color relies on a lengthy series of photoreceptor cell fate determination steps with one of the more important being the decision of which opsin(s) to express (5–7) and which ones to repress.

The *Drosophila* compound eye is composed of approximately 750 ommatidial-subunits containing eight photoreceptors each (8) (**Figure 1A**). Six photoreceptors (R1-R6) express *ninaE*-encoded Rhodopsin 1 (Rh1), are sensitive to both UV and blue light, and are largely used for achromatic vision and motion detection (similar to vertebrate rod photoreceptors) (9–13). Color vision is mediated by the remaining two photoreceptors, R7 and R8, which express some combination of ultraviolet-absorbing Rh3 or Rh4 (14–17), blue-absorbing Rh5 (18), or green-absorbing Rh6 (19–21). The majority of ommatidia in the retina can be classified as either pale (p) or yellow (y) and can be distinguished based on opsin expression (22). Rh3-expressing R7p cells are nearly always paired with an adjacent Rh5-expressing R8p, while Rh4 and Rh6 are expressed in R7y and R8y cells respectively (18, 23, 24). In addition to these predominant subtypes, a subset of R7y cells located in the dorsal 1/3rd region co-express Rh3 and Rh4 (25) while 1-2 rows of ommatidia on the dorsal periphery express Rh3 in both R7 and R8 photoreceptors (8, 26, 27).

**Figure 1:**
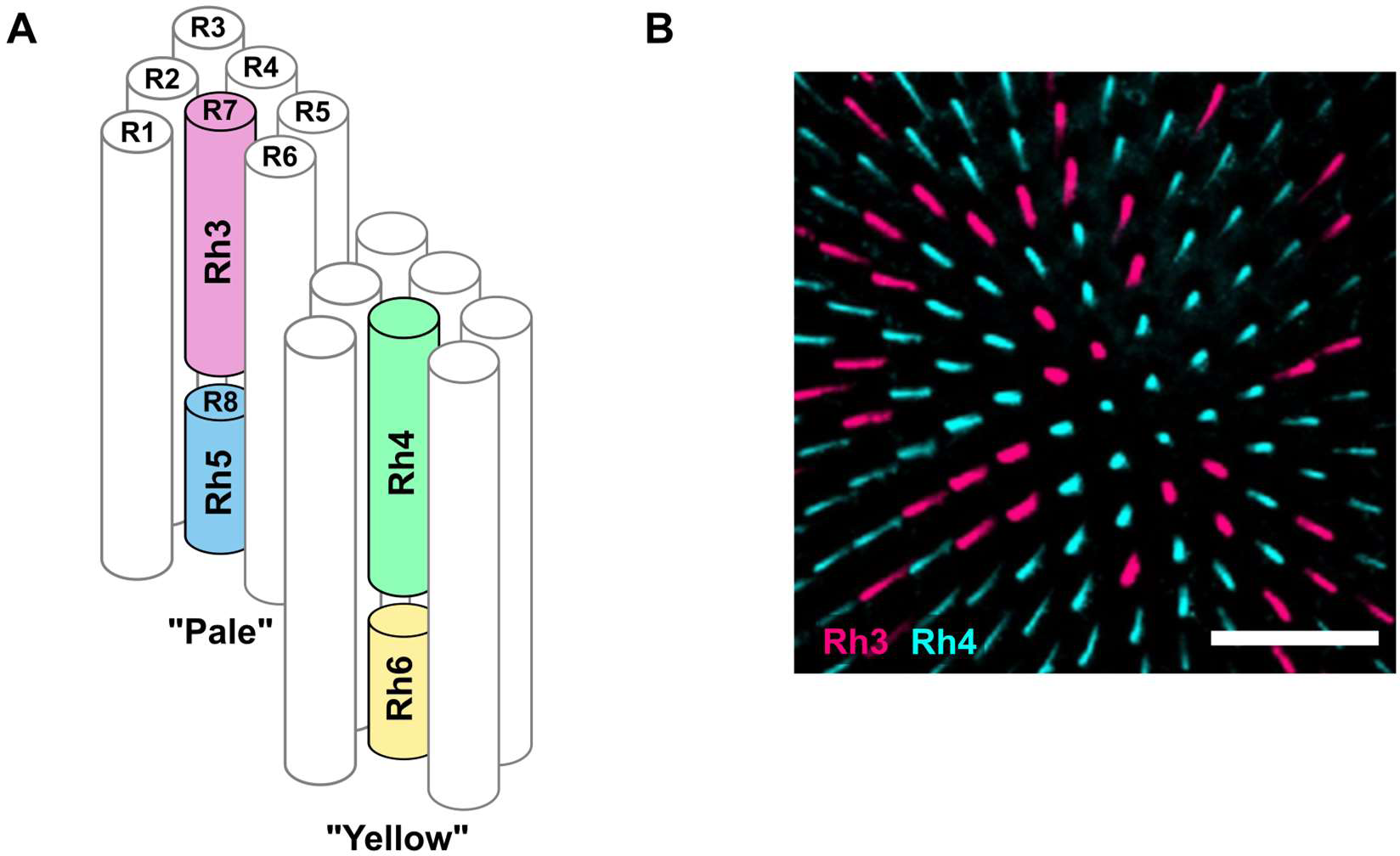
Photoreceptor subtypes are defined by stochastic and induced opsin expression. The *Drosophila* retina is composed of ∼750 photoreceptor clusters called ommatidia (**A**). Photoreceptors R1-6 express Rh1, the primary light-sensing molecule in *Drosophila* vision, while R7 and R8 mediate color vision and express various spectrally tuned, color-sensitive opsins. Opsins localize to specialized structures called rhabdomeres, which are represented as cylinders above. Most ommatidia are classified as either “pale” or “yellow” with the pale subtype coordinately expressing Rh3 and Rh5 in R7 and R8 respectively, while yellow express Rh4 and Rh6. The initial choice between Rh3 and Rh4 expression in R7 cells is a stochastic process that results in fairly consistent 3:7 pale:yellow ratio of R7 cells as shown in a whole mount retina from a *cn bw* control strain (**B**). In contrast, R8 opsin expression is based on the presence or absence of an inductive signal from the adjacent R7 cell. Scale bar represents 50 µm

The pale vs yellow fate decision in R7 and R8 cells is regulated by a combination of a stochastic and a deterministic process (28–30). After photoreceptor specification, at approximately 50% pupation, *spineless* (*ss*) is expressed in a subset of R7 photoreceptors where it mediates the activation of Rh4 and repression (via *defective proventriculus*) of Rh3 (31–34). *ss* is expressed in approximately 70% of R7 cells, which assume the yellow fate, while in the remaining 30% *ss* is repressed resulting in adoption of the Rh3-expressing pale identity (**Figure 1B**). Coordinate opsin expression is achieved in R8 cells via an inductive signal from the R7 (23, 24). In pale ommatidia, R7p instructs the adjacent R8 cell to express Rh5 and repress Rh6. In the absence of this signal, R8y cells express Rh6.

Many of the factors involved in regulating opsin expression in R8 cells or in establishing and/or maintaining the inductive signal from R7 to R8 have been identified in previous studies (35–42), while relatively few genes have been found that modulate the initial stochastic fate decision in the R7 (31, 38, 43, 44). Although the transcription factor/aryl-hydrocarbon receptor encoded by *ss* and its nuclear-translocating binding partner encoded by *tango* (*tgo*) have emerged as critical components of this process, the mechanism underlying stochastic *ss* expression in R7 cells remains unclear (38, 45). A zinc-finger transcription factor encoded by *klumpfuss* (*klu*) represses *ss* transcription and modulates the proportion of R7p:R7y cells in a dosage-dependent manner (46). *klu* itself is expressed in all developing R7 cells but at varying levels suggesting that binary (i.e., “on” vs. “off”) *ss* expression can be regulated by subtle variation in other factors. Furthermore, the observation that *spineless* remains “off” in over 20% of *klumpfuss*-null R7 cells raises the possibility that other factors may be involved (46).

In this study we examined R7 opsin expression in the *Drosophila* Genetic Reference Panel (DGRP), a collection of highly-inbred, fully-sequenced, genetically polymorphic, wild-caught strains (47, 48). We found that the proportion R7y:R7p photoreceptor cells is a highly variable genetic trait within the DGRP, suggesting that multiple genes likely influence the cell fate decision underlying R7 opsin expression. Using this information, we performed a Genome Wide Association Study (GWAS) and identified 42 significant Quantitative Trait Loci (QTL) associated with 28 unique genes. Further analysis using RNA-Seq data and genetic and physical interaction data reveals a network of candidate genes expressed in the developing retina that modulate stochastic cell fate decision and opsin choice in R7 photoreceptors.

## Results

R7 photoreceptors adopt either a pale (R7p) or yellow (R7y) identity based on a stochastic process involving *ss*-mediated activation of Rh4 and repression of Rh3 in R7y cells (5, 31). Although the canonical ratio of R7p to R7y cells is often expressed as roughly 3:7 (**Figure 1B**), deviation from this ratio has been induced via genetic manipulations (31, 32, 38, 43, 46). To better understand how natural genetic variation may influence this process, we used immunofluorescence microscopy to characterize Rh3 and Rh4 expression in 46 homozygous DGRP strains (**Figure 2; Table S1**) (47, 48). The percentage of R7p in the tested strains varied continuously between 25% and 61% with a mean of 41%, which is approximately 10% higher than our *cinnabar brown* (*cn bw*) laboratory control strain (42). After adjustments for the presence of various chromosomal inversions and *wolbachia* infection status (**Figure S1**)—we estimate the broad sense heritability of this trait as approximately 29%, which indicates the degree to which genotypic variation accounts for the observed variation in phenotype. Without phenotypic adjustment, we estimate broad sense heritability as 49%.

**Figure 2:**
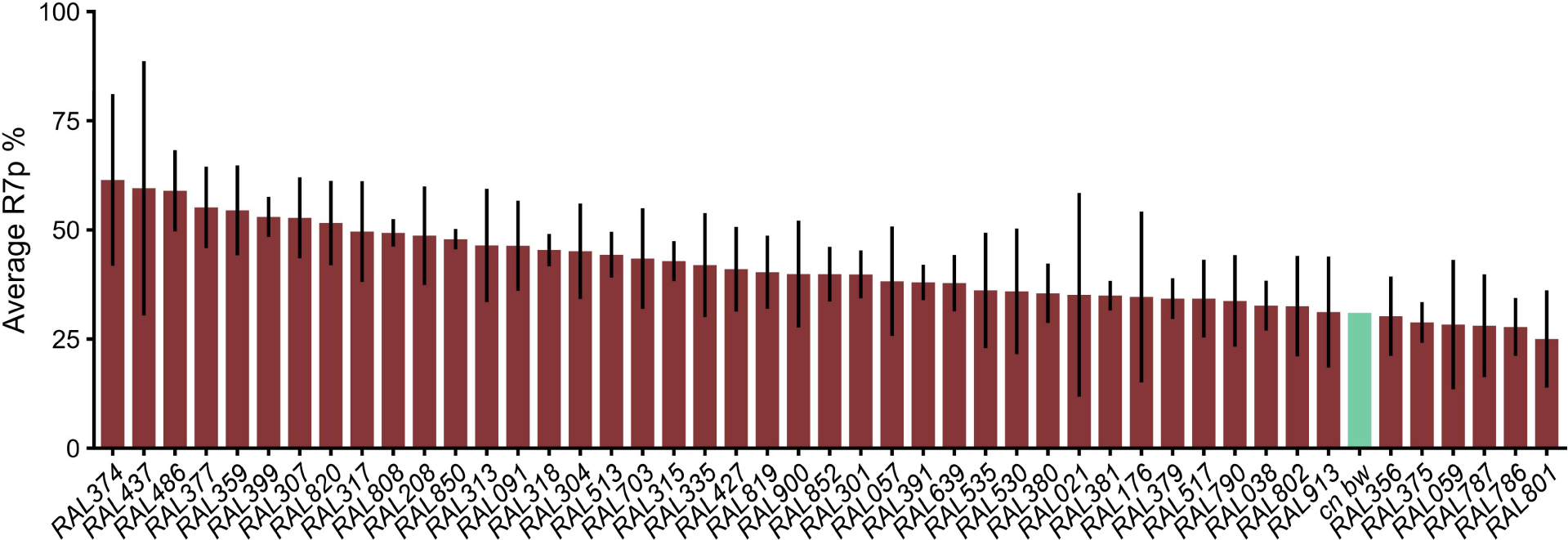
Natural genetic variation influences R7 subtype specification. The percentage of pale R7 cells (R7p) was determined in 46 DGRP strains based on Rh3 and Rh4 antibody staining (see **Methods**). Red bars represent line means, based on 3-8 replicate slides per strain while error bars represent +/- within-line standard deviation. The line mean from the *cn bw* laboratory strain was previously determined (43) and is included (in green) for comparison.

To identify specific genetic variants that influence R7 cell fate determination we performed a GWAS using the DGRP web tool (Freeze 2)(49), which adjusts for chromosomal inversions, *wolbachia* infection status, and population structure (48, 50). Overall, this analysis evaluated over 1.5 million variants with a minor allele frequency of at least 0.05 (**Figure 3**; **Table S2**). 42 variants were identified with a p-value lower than a cutoff of 1x10^-5^ and 26 of these mapped within 1 kb (upstream or downstream) of at least one annotated gene (**Table 1 & S3**). 12 of the significant variants mapped near multiple genes. For example, an SNP was identified on chromosome 2L that is located 27 bp upstream of *CG6144* on the plus strand and 45 bp upstream of *Cog4* on the minus strand—both genes were considered candidates for further analysis. In other cases, multiple significant variants mapped near the same gene—for instance, five separate SNPs map near candidate *CG31156*. 16 of the 42 variants are located further than 1 kb from an annotated gene. On average, the nearest gene to these “intergenic” variants is 7611 bp. Although there are many examples of cis-acting interactions at similar or even greater distances (46, 51)—to say nothing of inter-chromosomal interactions (52)—for this study, we chose to prioritize the 28 candidate genes located within 1 kb of at least one significant variant.

**Figure 3:**
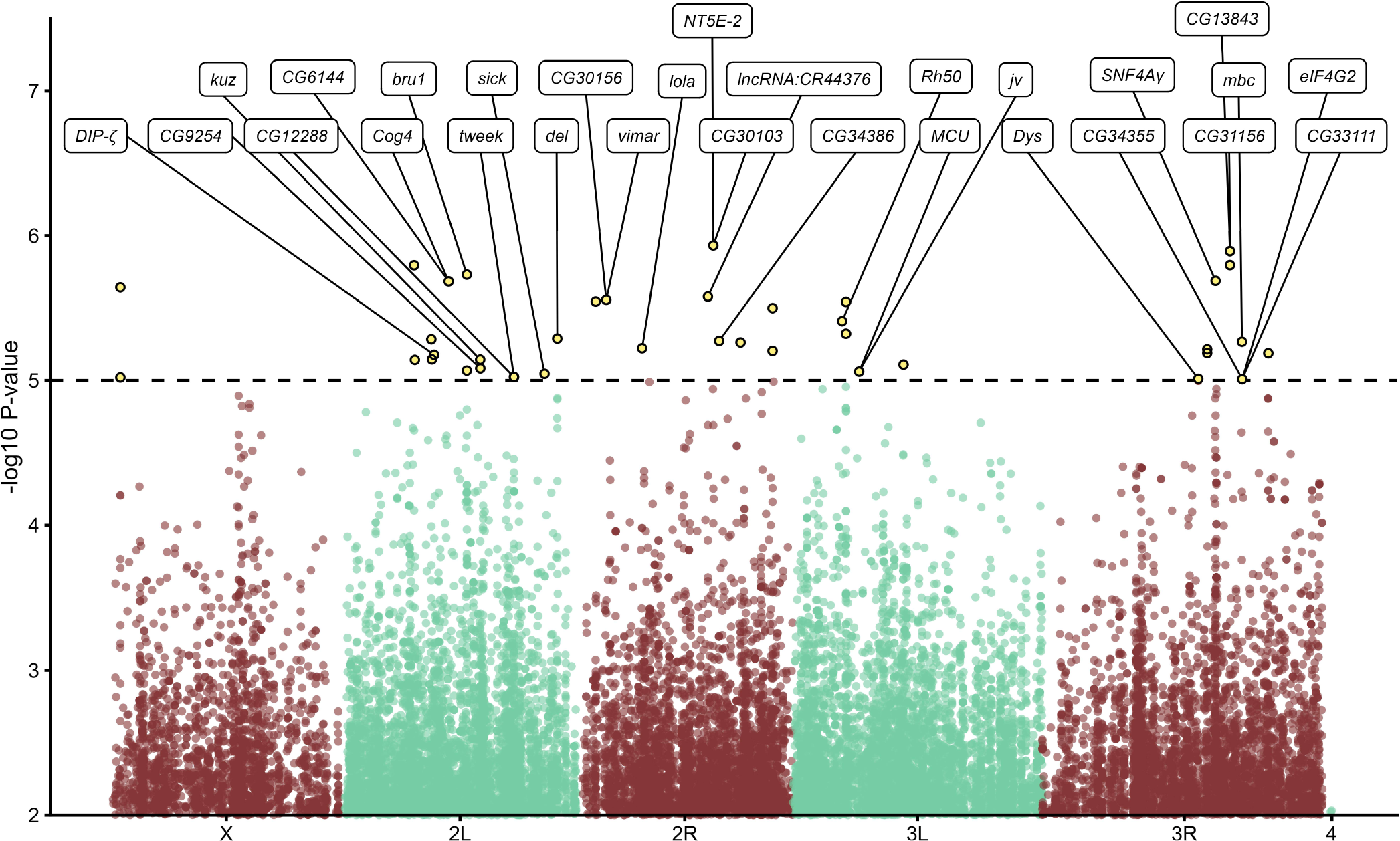
Genetic variants associated with % R7p variation. A GWAS was performed using the mean % R7p values shown in Figure 2, the results of which are summarized here. All genetic variants with p < 1x10^-2^ are plotted according to their relative genomic position with chromosome arms indicated on the x-axis. The significance threshold used for this study (p < 1x10^-5^) is indicated by the dashed line, with genetic variants above that line highlighted in yellow. Significant genetic variants that are within 1 kilobase of an annotated gene are labeled with the gene symbol. When more than 1 significant genetic variant is within 1 kilobase of a gene, only the variant with the smallest p-value (largest -log10 p-value) is labeled with the gene symbol.

**Table 1:**
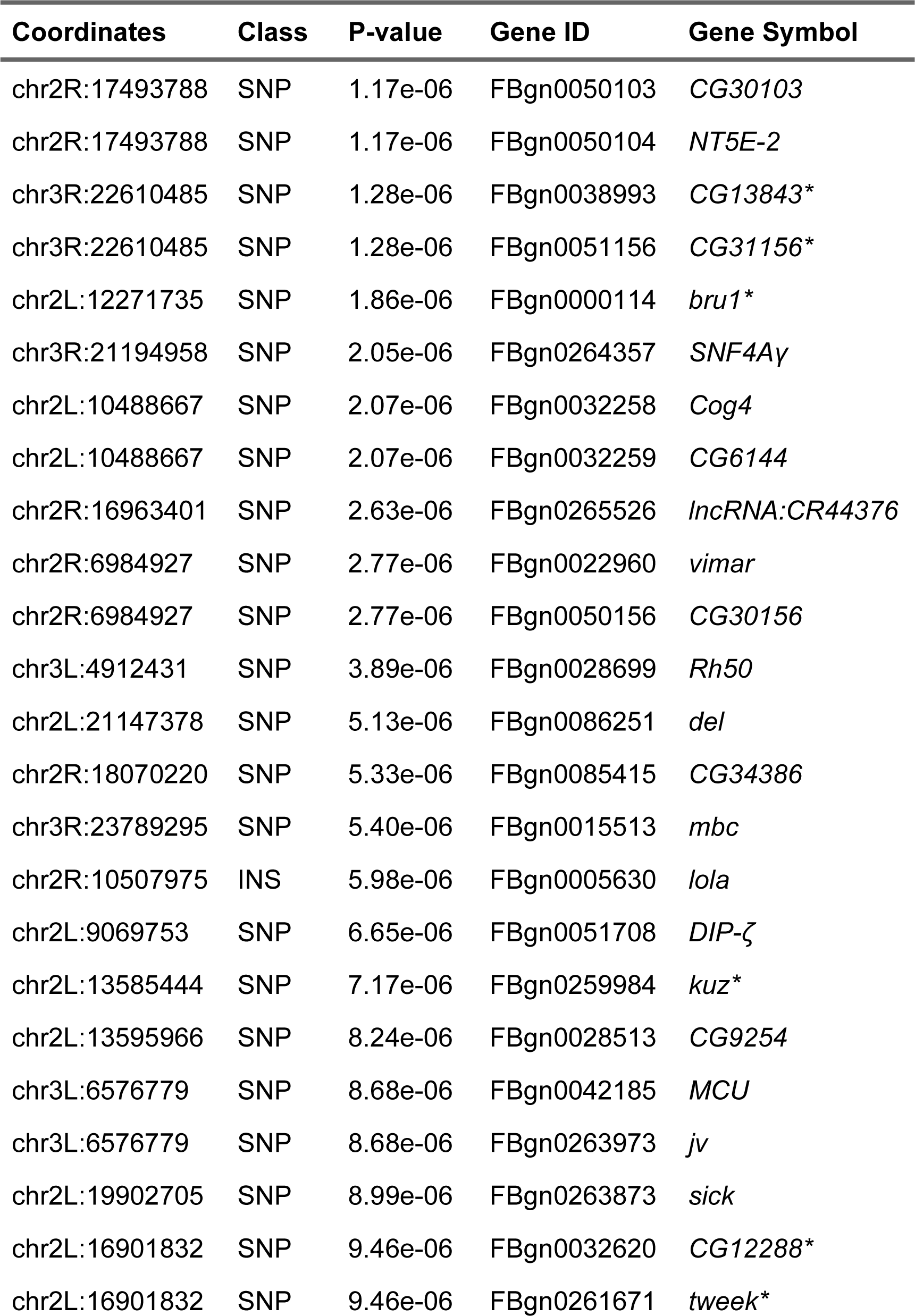
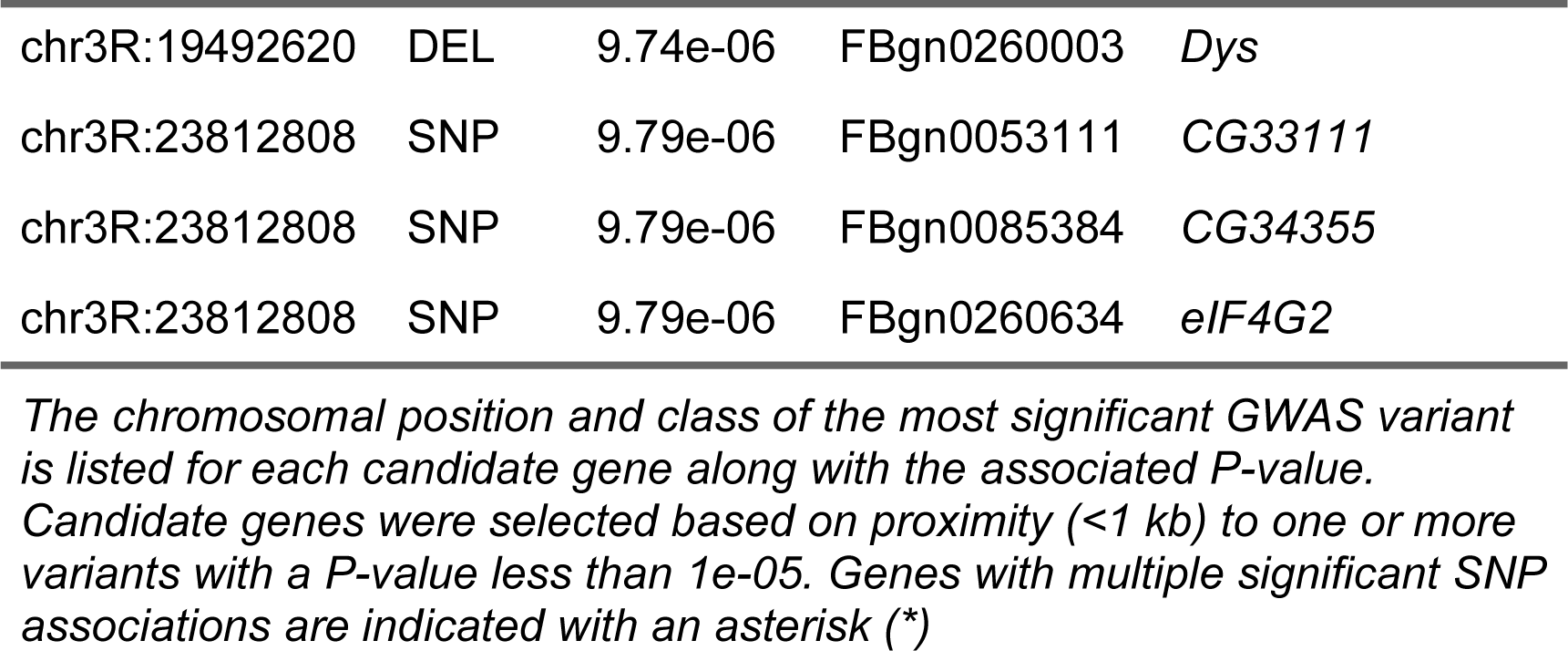
Candidate Modifiers of % R7p.

| **Coordinates** | **Class** | **P-value** | **Gene ID** | **Gene Symbol** |
| --- | --- | --- | --- | --- |
| chr2R:17493788 | SNP | 1.17e-06 | FBgn0050103 | *CG30103* |
| chr2R:17493788 | SNP | 1.17e-06 | FBgn0050104 | *NT5E-2* |
| chr3R:22610485 | SNP | 1.28e-06 | FBgn0038993 | *CG13843\** |
| chr3R:22610485 | SNP | 1.28e-06 | FBgn0051156 | *CG31156\** |
| chr2L:12271735 | SNP | 1.86e-06 | FBgn0000114 | *bru1\** |
| chr3R:21194958 | SNP | 2.05e-06 | FBgn0264357 | *SNF4Aγ* |
| chr2L:10488667 | SNP | 2.07e-06 | FBgn0032258 | *Cog4* |
| chr2L:10488667 | SNP | 2.07e-06 | FBgn0032259 | *CG6144* |
| chr2R:16963401 | SNP | 2.63e-06 | FBgn0265526 | *lncRNA:CR44376* |
| chr2R:6984927 | SNP | 2.77e-06 | FBgn0022960 | *vimar* |
| chr2R:6984927 | SNP | 2.77e-06 | FBgn0050156 | *CG30156* |
| chr3L:4912431 | SNP | 3.89e-06 | FBgn0028699 | *Rh50* |
| chr2L:21147378 | SNP | 5.13e-06 | FBgn0086251 | *del* |
| chr2R:18070220 | SNP | 5.33e-06 | FBgn0085415 | *CG34386* |
| chr3R:23789295 | SNP | 5.40e-06 | FBgn0015513 | *mbc* |
| chr2R:10507975 | INS | 5.98e-06 | FBgn0005630 | *lola* |
| chr2L:9069753 | SNP | 6.65e-06 | FBgn0051708 | *DIP-ζ* |
| chr2L:13585444 | SNP | 7.17e-06 | FBgn0259984 | *kuz\** |
| chr2L:13595966 | SNP | 8.24e-06 | FBgn0028513 | *CG9254* |
| chr3L:6576779 | SNP | 8.68e-06 | FBgn0042185 | *MCU* |
| chr3L:6576779 | SNP | 8.68e-06 | FBgn0263973 | *jv* |
| chr2L:19902705 | SNP | 8.99e-06 | FBgn0263873 | *sick* |
| chr2L:16901832 | SNP | 9.46e-06 | FBgn0032620 | *CG12288\** |
| chr2L:16901832 | SNP | 9.46e-06 | FBgn0261671 | *tweek\** |

**Table 1: Candidate Modifiers of % R7p**
| Coordinates | Class | P-value | Gene ID | Gene Symbol |
| --- | --- | --- | --- | --- |
| chr3R:19492620 | DEL | 9.74e-06 | FBgn0260003 | *Dys* |
| chr3R:23812808 | SNP | 9.79e-06 | FBgn0053111 | *CG33111* |
| chr3R:23812808 | SNP | 9.79e-06 | FBgn0085384 | *CG34355* |
| chr3R:23812808 | SNP | 9.79e-06 | FBgn0260634 | *eIF4G2* |

*The chromosomal position and class of the most significant GWAS variant is listed for each candidate gene along with the associated P-value. Candidate genes were selected based on proximity (<1 kb) to one or more variants with a P-value less than 1e-05. Genes with multiple significant SNP associations are indicated with an asterisk (\*)*

To define the relationship between the candidate genes identified via GWAS and the previously characterized mediators of R7 photoreceptor cell differentiation and opsin expression (*ss, tgo* and *klu*) we examined the known genetic and physical interactions between these genes collectively. 12 candidate genes are present in the esyN database, and nine of these have documented interactions with at least one other candidate either directly or via a single intermediary gene. Using this approach, we identified 22 additional genes that interact physically or genetically with two or more of the GWAS candidates identified in the screen, *ss, tgo* or *klu* (**Figure 4; Table S4**).

**Figure 4:**
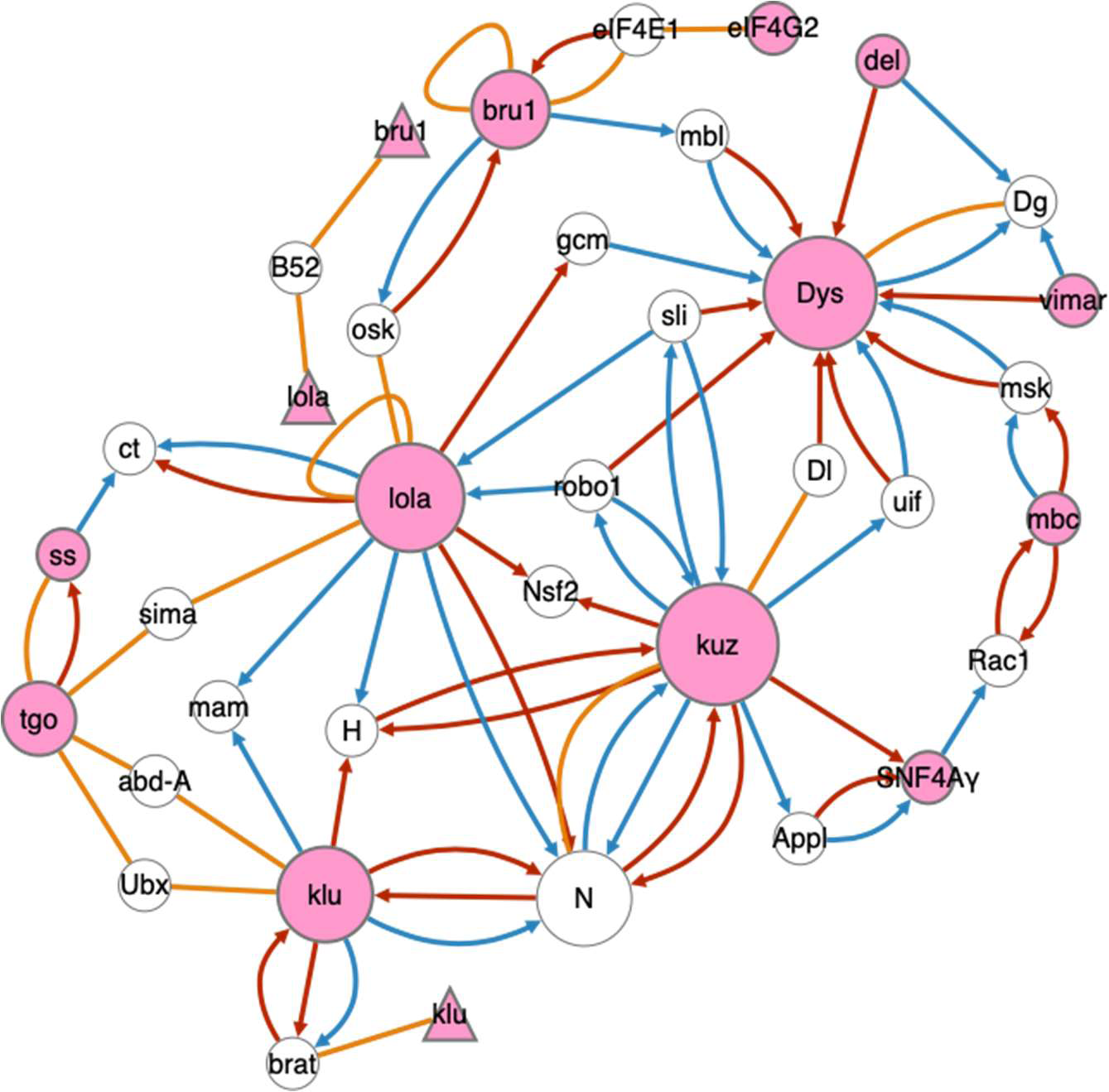
Network analysis of candidate genes. A network analysis was performed to identify physical and genetic interactions between candidate genes identified in the GWAS and the known regulators of Rh3 and Rh4 expression (*ss*, *tgo*, *klu*). The analysis was performed in esyN (53). Candidate genes are shaded in pink, physical interactions are shown in orange, genetic interactions are shown as arrows in the direction of the source gene enhancing (blue) or suppressing (red) the mutant phenotype of the target gene (using the nomenclature in FlyBase (54)). Genes or proteins are indicated as circles, while RNAs are shown as triangles. Additional interacting genes are included in the network (white circles) if they link two or more candidate genes.

Overall, these genes comprise a single large network where the genes *longitudinals lacking* (*lola), kuzbanian (kuz)* and *Dystrophin (Dys)* were the most highly connected (i.e., hub genes), each having connections to 9 or more other genes. This large network contains the known regulators of R7 photoreceptor cell specification *ss, tgo* and *klu*, and although they do not have direct genetic or physical interactions with any of the candidate genes we identified via GWAS, secondary interactions exist with *lola* and *kuz* through *Notch* (*N*), *Hairless* (*H*), *mastermind* (*mam*), *similar* (*sima*), and *cut* (*ct*). An additional small network consists of physical interactions between RNA transcripts of candidate genes *lola* and *bruno1* (*bru1*) and the protein B52. Overall, these interaction networks suggest numerous mechanisms for regulation of *ss*, *tgo* and *klu* by the genes identified in the screen.

As an *in vivo* validation of our GWAS findings, we used *GMR-GAL4* driven RNAi to knockdown expression of 20 candidate genes in the developing eye (55–57), with specific candidates being chosen based on RNAi line availability (58, 59). Of the 37 lines tested (some genes are represented by multiple lines), 12 had a significant reduction in the proportion of R7p compared to their respective controls, while three had morphological defects preventing accurate counts, which likely reflect the broad expression pattern of *GMR-GAL4* (**Figure 5; Table S5**) (56, 57, 60). The largest effects were seen with knockdown of *CG12288*, which is predicted to encode an RNA-binding protein orthologous to human RBM34, and *SNF4Aγ*, which encodes the Adenosine Monophosphate-binding subunit of the heterotrimeric AMP-activated Protein Kinase (61, 62).

**Figure 5:**
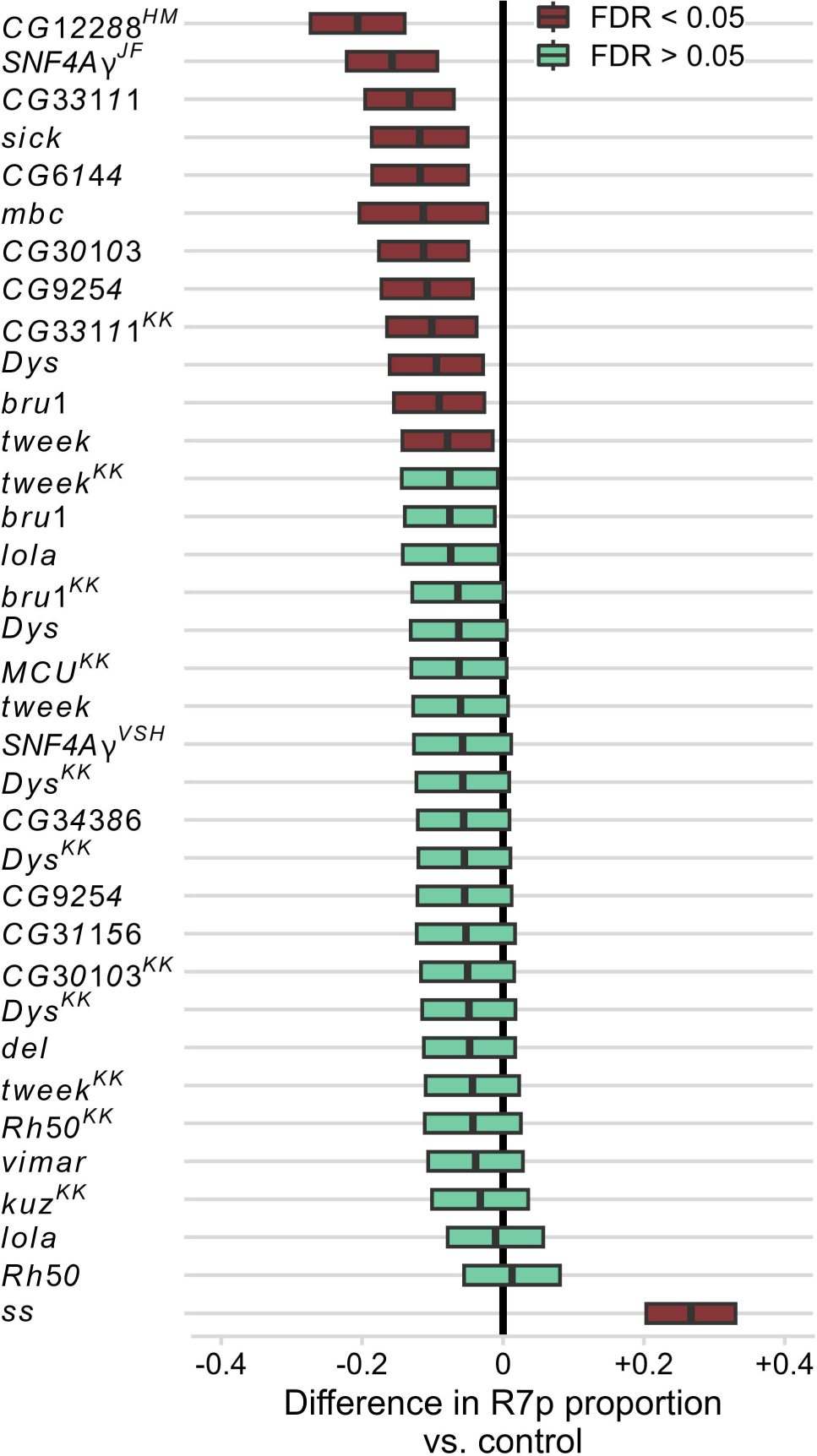
RNAi knockdown of candidate genetic modifiers of R7p proportion. *GMR-GAL4* was used to drive expression in the developing eye of one or more RNAi constructs targeting the indicated candidate genes (see **Table S5** for full details). The proportion of R7p was determined based on antibody staining of dissociated ommatida and is expressed as the difference from control animals. A vertical line intersecting “0” on the x-axis represents “no change” from control. The ends of the rectangle represented the upper and lower limits of the 95% confidence interval and the vertical line within the rectangle represents the difference in mean R7p proportion between the knockdown and the control. The color of the rectangle indicates statistical significance determined via a z-test (Red indicates FDR <0.05). Similarly-obtained values for *spineless* (*ss*) knockdown are included for comparison. The majority of RNAi lines are from the VDRC GD library (58), while a superscript denotes alternative VDRC stocks (KK and VSH) or TRiP (63) stocks (HM & JF)—see **Methods** for details.

To accompany our genomic approach, we used RNA-Seq to profile the developing retina transcriptome. Retinal tissue was collected at 40 hours after puparium formation (apf), which is after photoreceptor recruitment/specification has occurred (64–67), at the time when *spineless* is expressed (31), but before terminal differentiation and the completion of synaptogenesis (8, 68, 69). To identify transcripts specifically enriched in R7 photoreceptors, we collected samples from control animals (*w^1118^*) and *sevenless* loss of function mutants (*w* sev^14^*), which lack R7 cells (70, 71). After mapping and gene-level quantification, 13,517 genes (with TPM > 0 across all samples) were tested for differential expression using DESeq2 (see **Methods** for details). 917 genes in total were differentially expressed (FDR < 0.05) between control and *sev* mutant samples (**Figure 6** & **Table S6**). Of the 28 GWAS candidate genes, all but four were detected in developing retina at the critical time point when subtype identity is established, i.e., within the control samples (median TPM > 0; **Table S6**). Five are upregulated in *sev* mutant retinas (*NT5E-2, jv, CG31156, SNF4Aγ,* and *CG34386*) and none are significantly downregulated (**Figure 6A**), suggesting that—like *spineless,* which is heavily transcribed in bristle cells (31)—they are not necessarily enriched exclusively in R7 cells at this developmental stage.

**Figure 6:**
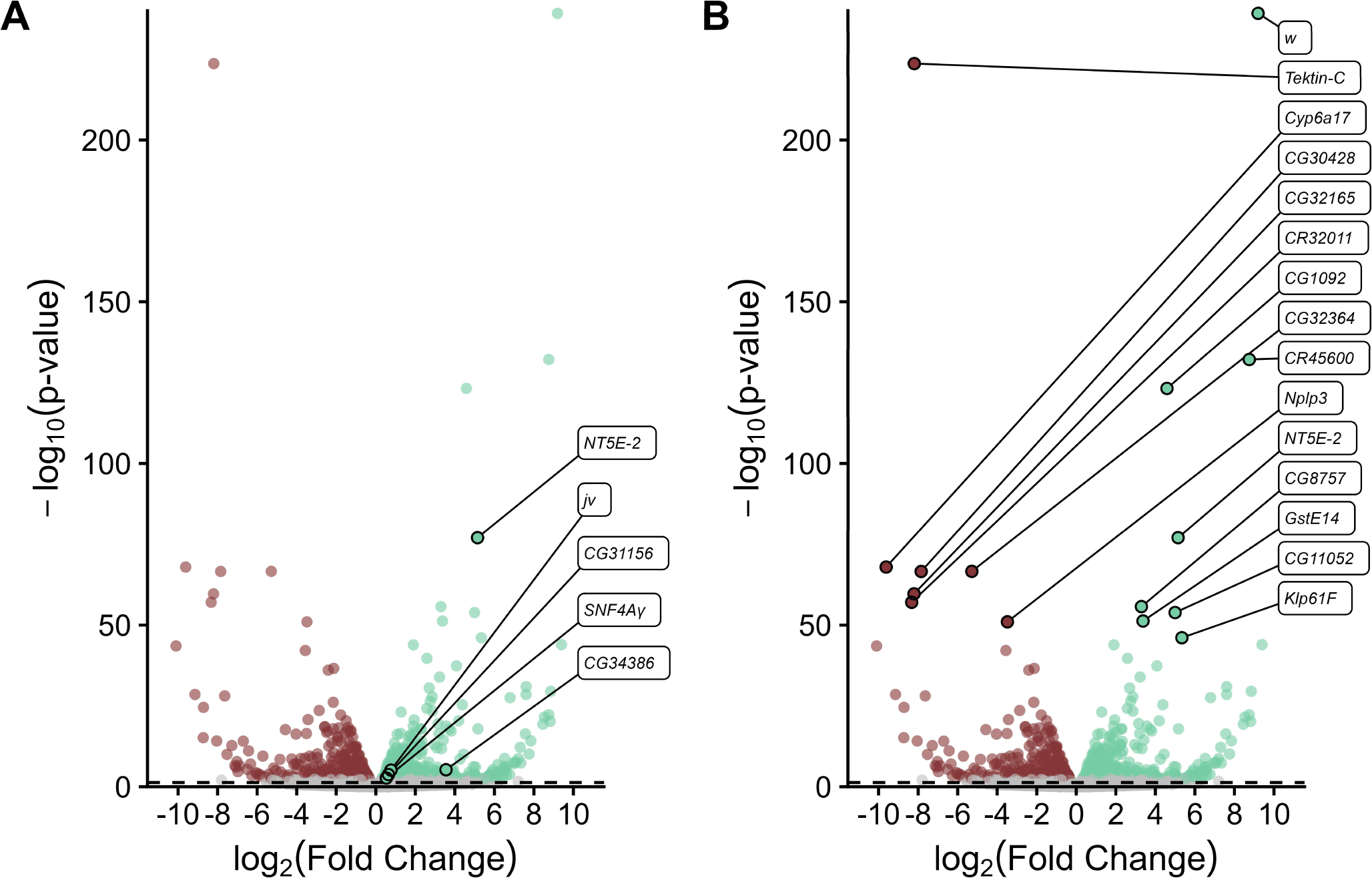
R7-dependent expression profile of the developing retina. RNA-Seq was performed using libraries generated from 40 hr apf retinas isolated from either *sevenless* (*sev*) (*w* sev^14^*) mutants, which lack R7 photoreceptors, or from control animals (*w^1118^*). The x-axis represents log_2_-transformed fold change (mutant vs. control) values for each gene while the y-axis shows log_10_-transformed FDR. Significantly (FDR < 0.05, indicated by dashed line) genes are highlighted in green while significantly downregulated genes are in red. Differentially expressed GWAS candidate genes are indicated in (**A**), which has a truncated y-axis to aid in separation of the data points, while (**B**) shows the full y-axis and has the top 15 differentially expressed genes labeled.

The 372 genes downregulated in *sev* mutants relative to the control (**Figure 6B** & **Table S6**) are particularly relevant to this study since we would expect that list to include genes that are highly enriched in the R7 photoreceptors during this critical time point and which may be important for opsin choice and terminal differentiation. In that regard, we note that in *sev* mutants, expression is largely eliminated (median TPM = 0) in 34 genes and reduced by at least half in 209. Downregulated genes may also include those associated with pale R8 photoreceptors—since most R8 cells adopt the yellow fate in the absence of R7 (24)—as well as some general photoreceptor-expressed factors due to the elimination of 1/8^th^ of the developing photoreceptors cells.

Conversely, we expect that the list of upregulated genes may include novel factors associated with yellow R8 photoreceptors as well as genes expressed in equatorial cone-cells—the cellular identity R7 cells adopt in the absence of *sevenless* (70, 71). We note that *white* is the top upregulated gene (**Figure 6B** & **Table S6**), which was unexpected since both the mutant and control strains are white-eyed and were thought to carry the transcriptionally-null partial deletion allele *w^1118^* (72, 73). Based on our RNA-Seq results, it is likely that the *sevenless* mutant stock we used carried an alternative allele that disrupts *white* function without eliminating transcription (74).

Finally, we identified and ranked statistically overrepresented Gene Ontology (GO) terms associated with our lists of upregulated and downregulated genes in *sev* mutants (**Table S7**) (75, 76). A number of GO terms related to chromatin regulation, e.g., "regulation of chromosome organization" (GO:0033044) and "positive regulation of heterochromatin formation" (GO:0031453), were enriched among the downregulated genes. In terms of specific genes, the suppressor of variegation genes *Su(var)3-7* and *Su(var)205* (aka *Heterochromatin Protein 1a*) stand out in particular due to their involvement in Position-effect variegation, a stochastic gene regulatory process phenomenon reminiscent of R7 cell fate specification (77, 78). A recent study illustrates this connection further by demonstrating the role of chromatin compaction dynamics in *spineless* expression (79). Analysis of upregulated genes reveals numerous GO terms related to chitin biosynthesis and/or cuticle development. We suspect this is most likely related to the transformation of R7 photoreceptors in *sev* mutants into lens secreting cone cells, which are cuticular in nature (70, 71, 80).

## Discussion

In a previous genetic screen (43), we identified a number of transposon insertions that could modulate R7 opsin choice either through ectopic misexpression or through gene disruption. These findings suggested that the relatively simple *ss*-mediated terminal cell fate decision in R7 photoreceptors may be influenced by a larger, more complex network of factors. In this study, we further explored this possibility by examining R7 opsin expression in a subset of the homozygous DGRP strains. We observed a wide degree of variability in this trait (**Figure 2**), much of which is likely attributable to the numerous genetic polymorphisms present in this collection. Further analysis via genome wide association identified 42 significant variants with the closest to *spineless* being over 3 Mb downstream.

In a prior GWAS study, Anderson and colleagues identified a *cis-*regulatory element bound by Klu approximately 7 kb upstream of *ss* that influences the proportion of R7p and R7y (46). This was achieved by outcrossing individual DGRP strains to a strain carrying a 197 kb *ss* deficiency (*Df(3R)Exel6269*) (81) in order to specifically enrich for *cis* elements affecting *ss* expression. Apart from the intended effect, this approach would likely reduce the effect of recessive QTLs while increasing the likelihood of detecting *ss*-interacting factors. After reanalyzing their phenotypic data via the standard DGRP pipeline (49), we found significant (p < 1x10^-5^) QTLs associated with *SNF4Aγ* and *Dystrophin (Dys)* (**Table S8**), two candidate genes identified in this study.

As part of the standard DGRP analysis pipeline, ANOVA is used to estimate the effects of *wolbachia* infection as well as the presence of five common chromosomal inversions. The submitted phenotype values are subsequently adjusted to account for the effects of these factors before being used for GWAS (47, 48, 82). We noticed in the ANOVA table generated by this analysis that one of the inversions, *In(3R)Mo*, is associated with an estimated 20% increase in R7p ommatidia (p = 0.0008509) (**Table S9**). *In(3R)Mo* is present in five of the strains (*RAL374, 437, 820, 317,* and *335*) we tested in this study, three of which are homozygous for the inversion (*RAL374, 437*, and *820*). Prior to adjustment, all five strains have above average % R7p, with the three homozygous strains ranking in the top 4 of all the tested strains (**Figure S3A**). We used Fast-LMM (83) to repeat the GWAS analysis with and without adjustment for *In(3R)Mo* and noted substantial differences in the results (**Figure S3B**; Pearson *r* = 0.646).

Looking specifically at SNPs on chromosome 3R we find that, without adjustment for *In(3R)Mo*, none of the genetic variants identified earlier in this study (**Figure 3**) reach the significance cutoff level (**Figure S3C**). In all cases, the inversion is co-inherited with the reference (i.e. “major”) allele—without adjustment, this significantly increases the % R7p value obscuring the relative differences in comparison to the alternative (i.e. “minor”) alleles (see **Figure S3D** for an example).

26 of the 42 significant variants identified in our GWAS are located within 1 kb of at least one annotated coding region. 18 of the candidates have been previously characterized in *Drosophila* and are known to be involved in a variety of diverse cellular and molecular processes, while eight of the uncharacterized genes have clear human orthologs (**Table 1 & S3**). Especially notable are the nine candidates with previously reported neuronal functions and/or phenotypes. For instance, *visceral mesodermal armadillo-repeats* (*vimar*) is involved in photoreceptor neurogenesis (84) along with the *Notch*-interacting factors *longitudinals lacking* (*lola)* and *kuzbanian (kuz)* (85–87).

*SNF4Aγ* and *Dys* alleles are both associated with various neurodegenerative phenotypes (88, 89), with *Dys* having an additional role in synapse function and synaptogenesis as do *Dpr-interacting protein zeta* (*DIP-ζ)* and *tweek* (90, 91). Finally, *javelin* (*jv)* encodes an actin associated protein required for mechanosensory bristle formation (92), while *sickie (sick)* and *lola* play a role in axon growth and guidance (93, 94). The genetic and physical interaction networks of the candidate genes reinforce the notion that the hub genes *lola*, *Dys* and *kuz* are excellent candidates for regulating *ss*, *tgo* and *klu.* The empirically determined genetic and physical interactions through *N, H, mam, sima,* and *ct* provide plausible mechanisms through which the candidate genes identified in the GWAS integrate with the canonical regulators of R7p vs. R7y photoreceptor specification and opsin expression.

Terminal differentiation of photoreceptors into the yellow or pale identity is initiated by a burst of *ss* expression in a subset of R7 cells during mid pupariation (31). Other factors influencing this process would presumably be expressed at a similar time and place. To explore this possibility, we used RNA-Seq to quantify mRNA expression in control and *sev* mutant pupal retinas (40 hr apf) (**Figure 6**). While all but four of the candidates were expressed to varying degrees, only five were differentially expressed in *sev* mutants. This suggests that even if these genes are expressed in R7 cells—which cannot be determined using this data—they are not expressed exclusively in R7s. Similarly, we find that *ss* is highly expressed in both control and mutant tissue, which is consistent with the prior observation that it is expressed in bristle cells of the developing retina in addition to a subset of R7 cells (31). Although none of the GWAS candidates were significantly reduced in *sev* mutant retinas, this analysis identified many genes that were (**Figure 6B** & **Table S6**). It is likely that among these genes are those that are either transcriptionally enriched in the R7 at 40 hr apf, or genes with expression that is dependent upon the presence of R7, such as a gene expressed in pale R8s.

To further evaluate our GWAS candidates we performed a secondary screen using 37 publicly available RNAi lines (58, 59). Although, 12 of the tested lines significantly reduced the R7p proportion compared to the control crosses, there was some amount of disagreement in cases where lines from different collections targeting the same gene were tested (**Figure 5**). For example, *GMR-GAL4* driven expression of the *UAS* construct *JF02060* to knockdown *SNF4Aγ* caused a significant reduction in R7p while *VSH330582* had no significant effect (**Table S5**). This discrepancy likely reflects one or more differing aspects of how the given RNAi strains were constructed including 1) the use of long double-stranded RNAs vs short, inverted repeats or small hairpins, 2) site-specific integration using ϕC31 or random P-element insertion, or 3) complimentary sequence selection within a target transcript.

The overall trend of R7p reduction, as opposed to increase, across the tested lines is also notable and suggests that there may be background effects present in the RNAi collections that are not fully reflected in the control phenotypes, or that R7p reduction may be the more favored outcome in this particular knockdown regime. R7p proportion is elevated in the RNAi controls compared to *cn bw* (∼50% vs 30%; **Table S5**), which may limit or obscure the degree of R7p increase / R7y decrease we are likely to observe. Indeed, *spineless* knockdown under these conditions produces a mild ∼25% increase in R7p, which, given its centrality to this process (31), may represent the upper limit of what can be achieved with these RNAi components.

Technical considerations aside, a similar trend was noted in a prior study using RNAi to validate GWAS candidates affecting visual senescence in *Drosophila* where the authors noted that the majority of *GMR-GAL4* driven knockdowns resulted in significantly reduced phototaxis (95). While polymorphisms may individually or collectively influence a quantitative trait—either positively or negatively—by affecting expression levels of components within of a critical pathway, severe reduction of gene expression via RNAi seems more likely to simply disrupt the entire network in a deleterious manner. In the context of R7 terminal differentiation, this suggests that an active repressive mechanism may be required for the stochastic *spineless* expression pattern underlying opsin choice. In such a model, *ss*, which is required for Rh4 expression and R7y fate specification, can be expressed in all R7 cells by default and is only silenced in a subset via an active stochastic mechanism. Disruption of this silencing mechanism either directly, by removing a critical component, or indirectly, by general R7 cellular perturbation, would result in an increase in R7y vs R7p cells.

Indeed, a pair of silencing elements flanking the *ss* coding sequence which are required for *ss* suppression have been previously identified (32), while a more recent study has described the formation of silent/closed chromatin at the *ss* locus in presumptive R7p cells (79).

## Conclusion

In this study we examined opsin expression in a collection of wild-caught, highly-inbred *Drosophila* strains and observed a wide degree of variation in the proportion of pale and yellow R7 cells suggesting that terminal differentiation in photoreceptors is influenced by naturally occurring genetic variation. Genome-wide association was used to identify 42 polymorphisms significantly associated with this trait along with 28 candidate genes (12 of which were validated using RNAi). Many of the candidate genes reside within a large genetic and physical interaction network and all but four are expressed in the developing retina at the critical time point when subtype identity is established.

Collectively, these results suggest that the relatively simple *ss*-mediated cell fate decision in R7 photoreceptors is modulated by a much larger network of factors.

## Methods

### Drosophila husbandry and genetics

All stocks and crosses were maintained in humidified incubators at 25°C on cornmeal media based on the Bloomington *Drosophila* Stock Center recipe (**Table 2**). DGRP strains (**Table 1**) are described here (47, 48) and were obtained from the Bloomington *Drosophila* Stock Center (NIH P40OD0185376) along with the *GMR-GAL4* driver line *w*; P{w^+mC^=GAL4-ninaE.GMR}12* (56) and various *UAS-RNAi* responder lines and controls (RRID:BDSC_35787 & RRID:BDSC_56037) generated by the Transgenic RNAi Project (59). Additional *UAS-RNAi* lines were obtained directly from the Vienna *Drosophila* Resource Center including background control stocks for GD, KK, and shRNA (VSH) stocks (VDRC # 60000, 60100, and 60200 respectively) (58). See **Table S5** for description of specific RNAi stocks used. In most cases, opsin expression was quantified (see below) in female progeny derived from female *GMR-GAL4* flies crossed to male *UAS-RNAi* or control flies. For TRiP RNAi lines, male progeny were used in order to avoid any confounding effect from the recessive *sev^21^* allele found in those stocks (96). Other strains used in this study include *w^1118^* (FBal0018186), *cn^1^ bw^1^* (FBst0000264; RRID:BDSC_264) and *w* sev^14^* (24, 72, 73).

**Table 2:** *Drosophila* Media Recipe.

| Ingredient | Amount | Manufacturer | Supplier | Catalog # |
| --- | --- | --- | --- | --- |
| Water | 39L | NA | NA | NA |
| Inactive Dry Yeast | 675g | Flystuff | Genesee Scientific | 62-106 |
| Soy Flour | 390g | NutriSoy | Genesee Scientific | 62-115 |
| Yellow cornmeal | 2850g | Quaker | Genesee Scientific | 62-101 |
| Light malt extract | 1800g | Flystuff | Genesee Scientific | 62-110 |
| Agar | 225g | Flystuff | Genesee Scientific | 66-103 |
| Light corn syrup | 3L | Karo | Various | NA |
| Propionic acid | 188mL | Fisher Chemical | Fisher Scientific | A258-500 |

*Recipe based on Bloomington Drosophila Stock Center's standard corn media. Makes approximately 47.5L. Propionic acid is added once the media has cooled just prior to dispensing.*

### Immunofluorescence microscopy

R7 subtype quantification was performed on dissociated ommatidia preparations as previously described (18, 97). Briefly, adult eyes were dissected from isolated heads (20 eyes per sample unless otherwise noted) in 100 µL 1X Phosphate Buffered Saline (PBS) using either a 28G needle or microdissection scissors (F.S.T. 15000-08). Isolated eyes were then sheared into smaller pieces before being triturated 10 times with a P200 micropipette with attached 200 μL tip. Samples were then transferred to a 25 X 75 X 1 mm Superfrost microscope slide (Fisherbrand 12550143) and allowed to dry on a 45°C slide warmer before being stored at -20°C. To facilitate whole slide imaging for the RNAi validation experiments (see below) triturated samples were passed through a 100 μm filter (CellTrics 04-0042-2318) before being transferred to the slide.

The immunostaining procedure was performed as previously described (18). Frozen slides were allowed to reach room temperature before being rehydrated in 1X PBS and dried on the slide warmer; all subsequent steps were carried out at room temperature. Samples were fixed for 10 minutes in 3% paraformaldehyde prepared in 1X PBS and then washed (5 min per wash) twice with 1X PBS, once with cytoskeletal buffer (10mM Hepes pH 7.4, 100 mM sucrose, 3 mM MgCl_2_, 50 mM NaCl, 0.5% Triton X-100, and 0.02% NaN_3_), and once more with 1X PBS containing 0.01% saponin.

Antibodies (see below for details) were diluted in 1X PBS containing 3% Normal Goat Serum, 1 mg/ml BSA, and 0.03% Triton X-100 and applied directly to the slides for 1 hour. Slides were then washed three times in 1X PBS containing 0.01% saponin before being mounted in ∼12.5 µL PermaFluor (ThermoFisher TA-006-FM), covered with a #1.5 coverslip, and sealed with clear nail polish. In instances where indirect immunofluorescence was used, three additional washes were performed in between primary and secondary antibody applications.

Rh3 was detected directly using an Alexa Fluor 647-conjugated mouse monoclonal antibody diluted 1:100 (98). Rh4 was detected either directly using an Alexa Fluor 488-conjugated mouse monoclonal antibody (24) diluted 1:100 or indirectly using a rabbit polyclonal antibody (99) (1:10 dilution) in conjunction with a goat anti-rabbit conjugated secondary (Jackson ImmunoResearch 111-295-003 or Invitrogen A11008) diluted 1:1000. For RNAi validation experiments R8s were counterstained using a combination of mouse monoclonal anti-Rh5 and anti-Rh6 (24), both conjugated with Alexa Fluor 555 (1:100). Antibody conjugations were performed using Alex Fluor protein/antibody labeling kits from Invitrogen (A20173, S10900, and A20187).

Opsin expression in the DGRP strains was quantified by direct visualization on an Axioskop plus (Carl Zeiss, Inc., Thornwood, NY) as previously described (43). For RNAi validation experiments, we used a Nikon Ni-E epifluorescence microscope (provided by the UT CBRS Microscopy and Imaging Facility) equipped with a motorized stage along with Nikon’s Elements imaging software (AR 5.21.02) to scan and merge 100 20X images covering ∼1 cm^2^ from each slide. R7p vs R7y was then manually scored in ImageJ v2.1.0 Fiji distribution (100, 101) using the cell counter plugin. In both cases, R7 cells expressing Rh3 exclusively were scored as pale, while those expressing Rh4 were scored as yellow.

The whole mount retina image in **Figure 1** is a maximum intensity projection reconstructed in ImageJ from a series of thirteen 1.5 µm z-stacks acquired on a Nikon A1R Confocal microscope. The dissection and immunostaining procedure was adapted from (102, 103). Whole eyes were dissected in 1X PBS supplemented with 0.1% Triton X-100 (PBT) using forceps and microdissection scissors. Eyes were then fixed for 1.5 hours at room temperature in 4% paraformaldehyde prepared in 1X PBS before being washed 3 times in PBT (30 min each) at room temperature. Eyes were incubated overnight with conjugated antibodies (described above), washed three more times (10 min each), and mounted in PermaFluor on a bridge slide with the cornea/lens facing the coverslip.

### Quantitative genetics and Genome Wide Association

Average R7p % was calculated from 3-8 replicate slides for each DGRP line tested. Broad based heritability was estimated as the coefficient of determination (R^2^) from a 1- way ANOVA using strain ID to predict R7p %. GWAS was performed using the DGRP web tool (49), which is described here (47, 48) and is based on the FaST linear mixed models algorithm (83). Although there are 4.4 million markers in the DGRP our analysis is limited to those with a minor allele frequency of at least 5% for the 46 strains examined. The “single mixed P-value” generated by this pipeline adjusts for population structure, the presence of various chromosomal inversions, and *wolbachia* infection status. QTLs with a single mixed P-value less than 1x10^-5^ were considered significant for this study. Candidate genes located within 1 kb from a significant QTL were chosen for further evaluation. To facilitate comparison with the RNA-Seq data, variant coordinates were updated from release 5 of the *Drosophila* genome to release 6 (BDGP6 aka dm6) using the LiftOver tool from the UCSC Genome Browser (104, 105). Similarly, gene annotations were updated to the Ensembl (release 89) gene build based on FlyBase release 6.02 (FB2014_05) using BEDOPS v2.4.29 (54, 106, 107).

Candidate gene information (name, symbol, summary) was taken from FlyBase release 6.38 (FB2021_01). To compare the effect of *In(3R)Mo* adjustment (**Figure S3**), we performed a GWAS using Fast-LMM (83), using SNPs with a MAF of at least 5%, and a pre-computed DGRP relationship matrix (82) in an effort to emulate the DGRP2 pipeline as closely as possible.

### Network Analysis

esyN (version 2.1) (53) was used to construct networks involved in R7 cell fate and opsin choice by extracting known genetic and physical interactions among candidate genes from the GWAS and *tgo, ss and klu,* genes known to be involved in R7 cell fate specification (based on FlyBase, version FB2022_03). Networks were restricted to include only connections that involved at least two candidate genes.

### RNA-Sequencing and differential gene expression analysis

White pre-pupae were collected and aged 40 hours at 25°C. Retinas were dissected (102, 108) in cold 1X PBS and transferred to a microcentrifuge tube containing RNAlater (Invitrogen, AM7020). Four replicate samples consisting of 10-20 retinas were prepared for each genotype (*w^1118^* or *w^1^ sev^14^*). Total RNA was isolated using the PureLink RNA purification kit (Invitrogen, 12183020) according to the manufacturer’s instructions along with on-column PureLink DNase treatment (Invitrogen, 12185010).

Library construction (from polyA-selected RNA) and sequencing was performed by the Genomic Sequencing and Analysis Facility at UT Austin, Center for Biomedical Research Support (RRID# SCR_021713) using the Illumnia MiSeq v3 Reagent Kit (paired-end, 150 cycles; MS-102-3001) and platform.

Raw paired-end reads were examined with FastQC (v0.11.4) (109) and adapters were trimmed with cutadapt (v1.9.1) (110) resulting in an average read length of 135 nt. The trimmed reads were then aligned to the Ensembl (release 89) transcriptome and genes were quantified with RSEM (v1.2.31) (107, 111) (**Table S10**). Transcripts with zero counts across all samples were filtered out of the dataset before using DESeq2 (v3.6) (112) to identify genes that are differentially expressed between the two genotypes. The Benjamini-Hochberg False Discovery Rate (FDR) was used to correct for multiple testing and significance was set at FDR < 0.05. (113).

GO term overrepresentation analysis was performed using the PANTHER statistical overrepresentation test found at http://www.pantherdb.org (Release 17.0) (75, 76). Analysis was performed separately on lists of upregulated and downregulated gene IDs with a reference list comprised of all the genes from the filtered list described above. Only the Complete GO annotation datasets for biological process, molecular function, and cellular component were evaluated.

### Additional data analysis

Statistical significance in the RNAi validation experiment was determined via a two-sided z-test comparing R7p proportions from test and control crosses—for this, we used the *prop.test* function from the stats package in R (v4.0.3) along with the *p.adjust* function for FDR correction (113, 114). Additional data analysis and visualization was performed in RStudio (v1.2.5033) (115) using the following packages: dplyr (v1.0.5), splitstackshape (v1.4.8), tidyr (v1.1.3), broom (v0.7.6), data.table (v1.14.0), ggplot2 (v3.3.5), cowplot (v1.1.1), viridis (v0.6.0), officer (v0.3.18), and flextable (v0.6.5). **Figure 1** was assembled using Inkscape (v1.0.1).

## Data availability

Raw paired-end reads as well as processed DESeq2 results are deposited under accession number GSE220550 in the Gene Expression Omnibus.

## List of abbreviations

GWAS: Genome Wide Association Study
QTL: Quantitative Trait Loci
R1-8: Photoreceptor cells 1 through 8
R7p: Pale R7 photoreceptors
R7y: Yellow R7 photoreceptors
Rh1-6: Rhodopsin 1 through 6
*ss*: *spineless* gene
Ss: Spineless protein encoded by *ss*
FDR: False Discovery Rate
kb: kilobase pairs
Mb: megabase pairs

## Acknowledgements

We thank Brenna J.C. Dennison, Meridee P. Mannino and Betelehem W. Yacob for assistance with sample preparation and data collection. This work was supported by grant R21EY024413 (SGB) from the National Institutes of Health.

## Supplementary Information

**Supplemental Figure S1:**
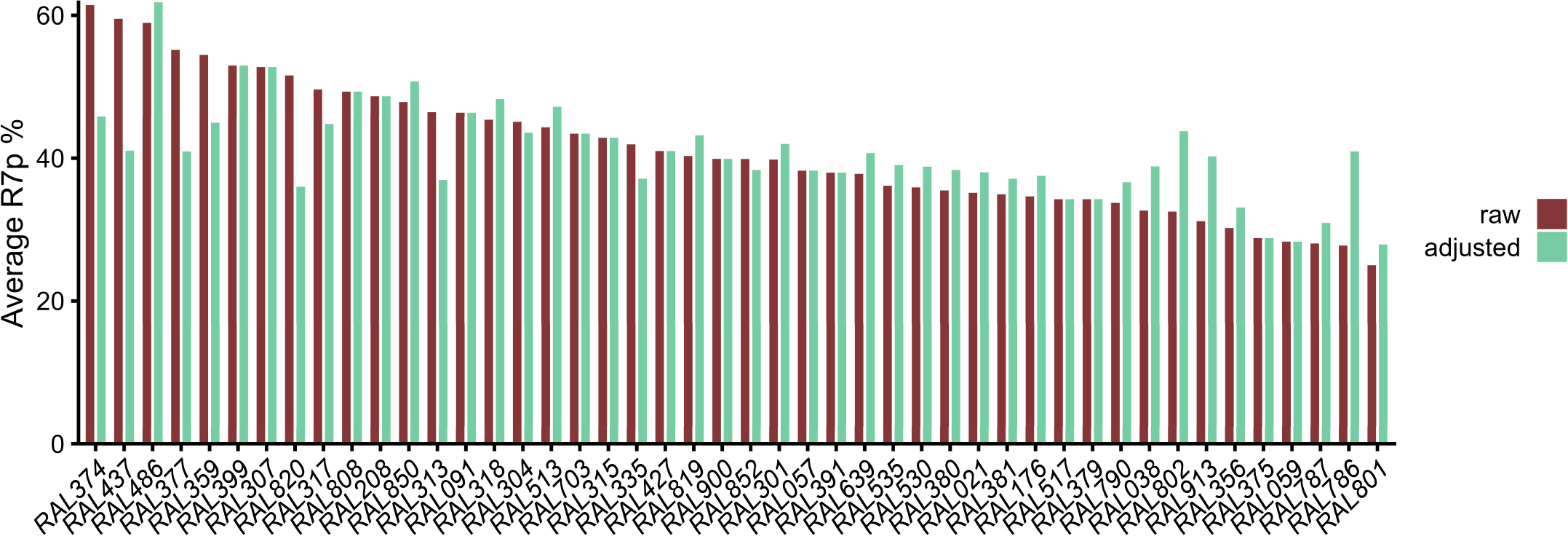
Average R7 pale percentage value is shown for the 46 tested DGRP strains in red, as in Figure 2. Adjusted phenotypes—compensating for *wolbachia* infection status and presence of various common chromosomal inversions within the DGRP—are shown in green.

**Supplemental Figure S2:**
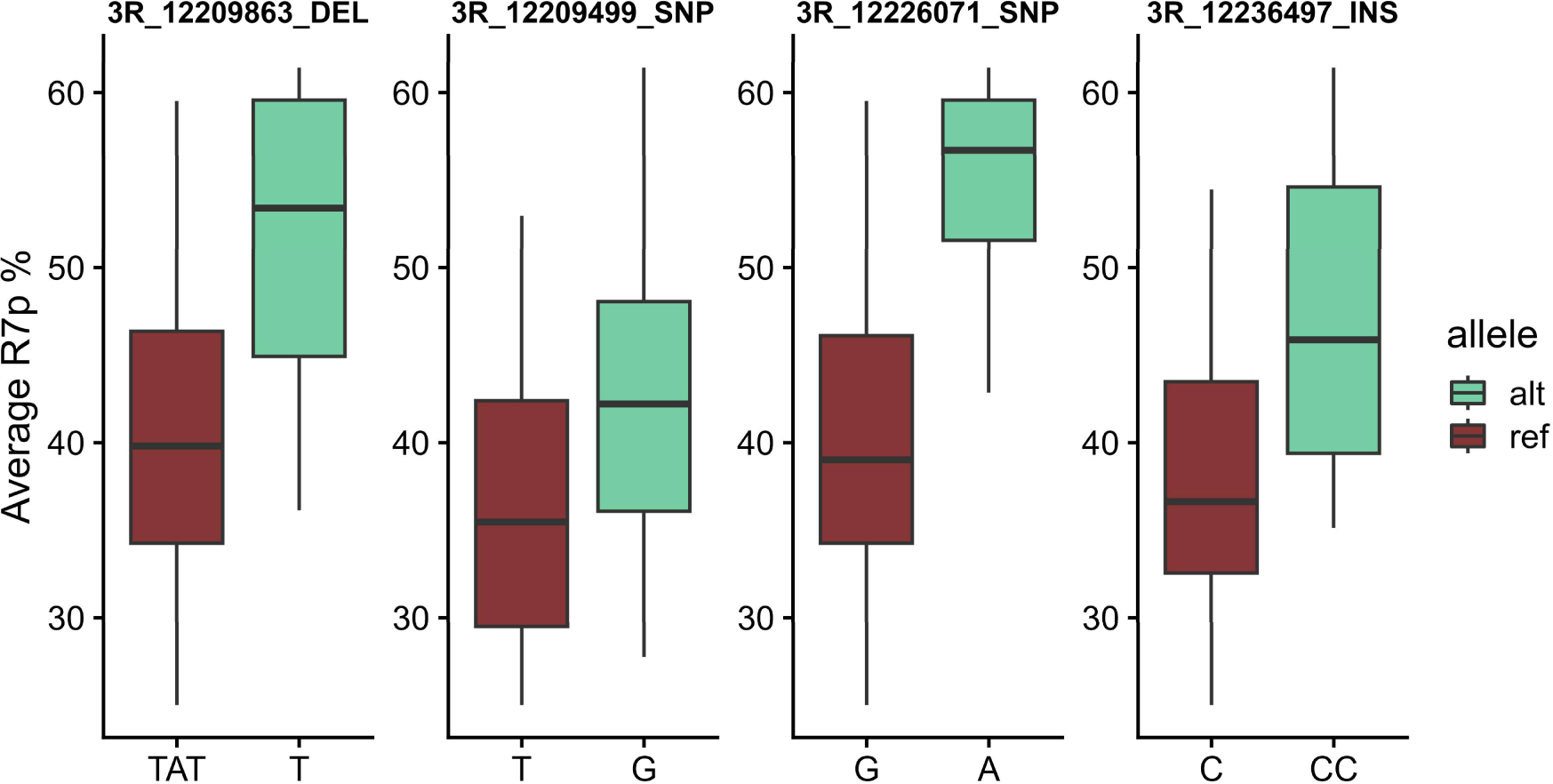
Box and whisker plot showing the average R7p% for DGRP strains with the indicated genotype for the top three (based on p-value) variants associated with *spineless*—i.e. within kb upstream or downstream of the coding sequence—along with the previously described (46) *ss^sin^* insertion (3R_12236497_INS). Values from strains homozygous for the reference allele (ref) are shown in red while those from strains homozygous for the alternative allele (alt) are shown in green.

**Supplemental Figure S3:**
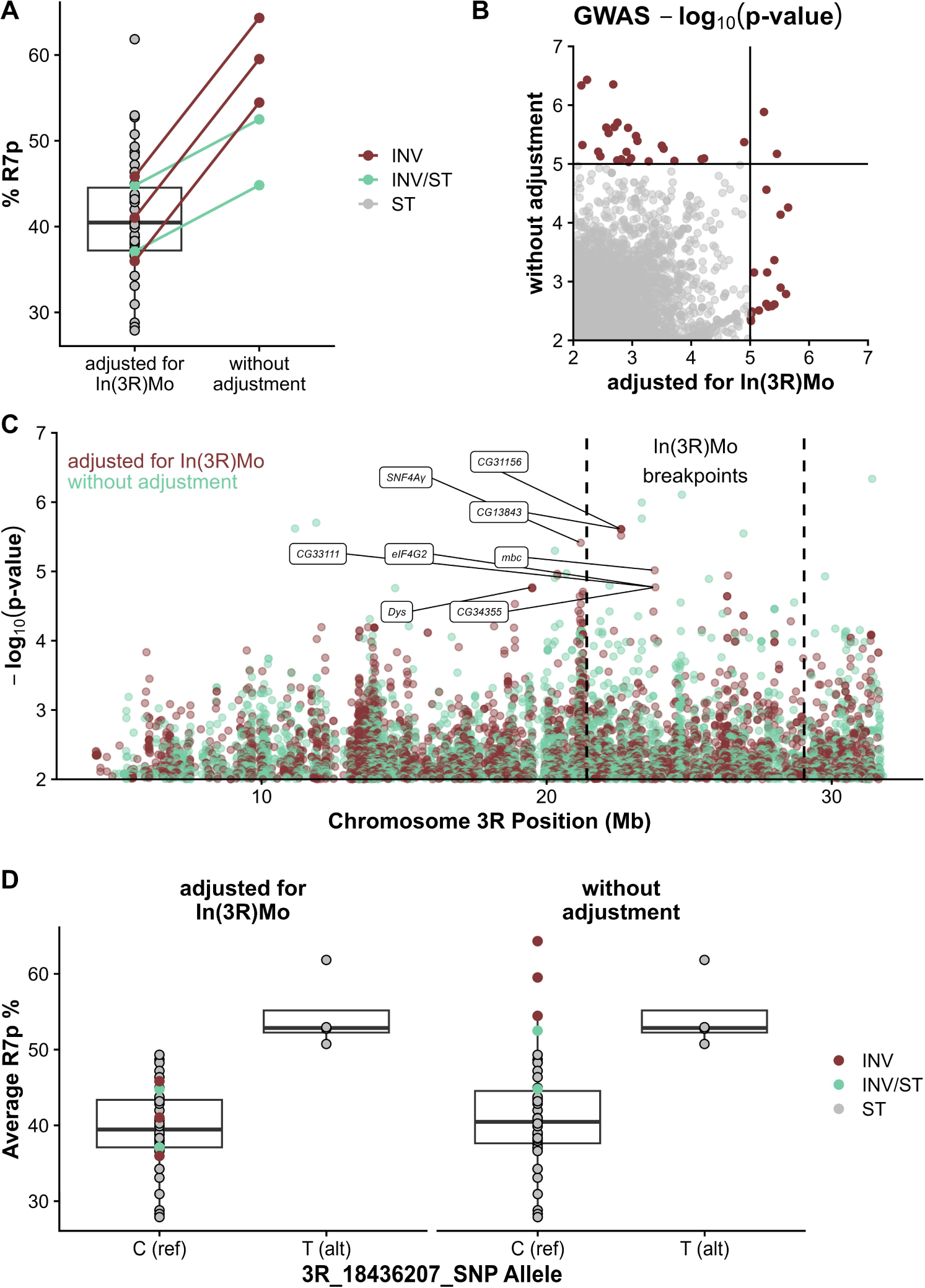
R7p % values adjusted for *Wolbachia* infection status and the presence of five major chromosomal inversions are shown in the left box plot (**A**). Five strains are either homozygous (INV; red) or heterozygous (INV/ST; green) for the *In(3R)Mo* inversion, while the rest have the standard (ST; grey) orientation of chromosome 3R. RYp % values without adjustment for *In(3R)Mo* are shown on the right—adjustments for *Wolbachia* and the other four inversions have still performed. GWAS was performed using R7p % values adjusted and unadjusted for *In(3R)Mo*, with subset of the results displayed in (**B**) as -log_10_ transformed p-values. Horizontal and vertical lines indicate the p-value cutoff of 1x10^-5^ with significant SNPs indicated in red. SNP -log_10_ transformed p-values from the *In(3R)Mo* adjusted (red) and unadjusted (green) datasets are plotted according to their position along chromosome 3R in (**C**) with the breakpoints of the inversion indicated by dashed lines. As in Figure 3, Significant SNPs that are within 1 kb of an annotated gene are labeled with the gene symbol. When more than 1 significant genetic variant is within 1 kb of a gene, only the variant with the smallest p-value (largest -log10 p-value) is labeled. Boxplots in (**D**) show % R7p for strains with either the reverence (ref) or alternative (alt) allele of SNP 3R_18436207. *In(3R)Mo* genotype is indicated as in (**A**) to illustrate the effect of adjustment on the left or no adjustment on the right.

**Supplemental Table S1**

Average R7 pale percentage values for the 46 tested DGRP strains along with the number of replicate slides counted per strain, standard deviation of replicate slides (SD), and the average R7 pale percentage values adjusted for *wolbachia* infection status and presence of various common chromosomal inversions within the DGRP.

**Supplemental Table S2**

GWAS data obtained via the DGRP2 analysis pipeline (49). The genomic coordinates of each tested variant are listed along with dm6-updated coordinates. P-values were adjusted via the linear mixed model approach to account for population structure and other forms of cryptic relatedness (83). Gene IDs are provided for any gene located within 1 kb of a variant.

**Supplemental Table S3**

Additional candidate gene information (full name & gene snapshot) obtained from FlyBase release 6.38 (FB2021_01).

**Supplemental Table S4**

Summary of network analysis. The table includes the candidate genes and additional interacting genes shown in Figure 4. For each interaction, the table indicates the source gene, type of interaction, the target gene and PubMed citation(s) (PMID) for the interaction. For genetic interactions, the table format indicates that the mutant phenotype of the target gene is suppressible (enhanceable) by a mutation in the source gene.

**Supplemental Table S5**

Stock information and R7 pale proportion values for tested RNAi lines. Males from the indicated RNAi stocks were crossed to female *GMR-GAL4* flies and opsin expression was quantified in progeny of the indicated sex. The prop.test function from the stats package in R (v4.0.3) along with the p.adjust function for FDR correction was used to test for significant difference in R7p proportion vs control flies generated by crossing *GMR-GAL4* females to appropriate background control strains (see **Methods** for details) (113, 114).

**Supplemental Table S6**

Summary of RNA-Seq results. Median normalized counts as well as rlog-transformed values are provided for all genes with counts > 0 across any sample. DESeq2 (112) was used to test for differential expression between *sev* and control tissues with the resulting P-values being corrected for multiple testing via the Benjamini-Hochberg method (113).

**Supplemental Table S7**

Results from the PANTHER statistical overrepresentation test found at http://www.pantherdb.org (Release 17.0) (75, 76). GO terms from the biological process, molecular function, and cellular component annotation datasets that are statistically overrepresented (p < 0.05) in the list of genes upregulated or downregulated in *sevenless* mutant retinas (**Table S6**) are given along with the gene IDs for the differentially expressed genes associated with that term.

**Supplemental Table S8**

GWAS data generated via the DGRP2 analysis pipeline (49) based on opsin expression values obtained from Anderson et. al, 2017 (46). The genomic coordinates of each tested variant are listed along with dm6-updated coordinates. P-values were adjusted via the linear mixed model approach to account for population structure and other forms of cryptic relatedness (83)—only variants with p < 1x10^-5^ are included. Gene IDs are provided for any gene located within 1 kb of a variant.

**Supplemental Table S9**

Type III ANOVA table showing the effects of five major chromosomal inversions and Wolbachia infection status on R7p %. Degrees of freedom (Df); Sum of Squares (sq); residual sum of squares (RSS); Akaike information criterion (AIC); F value and p-value (Pr (>F)). The estimated effects of each factor are given in the lower table along with standard error, t-value, and p-value.

**Supplemental Table S10**

Alignment statistics for RNA-Seq replicate samples.

